# Divergence in bacterial ecology is reflected by difference in population genetic structure, phage-predator load and host range

**DOI:** 10.1101/2022.09.06.506642

**Authors:** Karine Cahier, Damien Piel, Rubén Barcia-Cruz, David Goudenège, K. Mathias Wegner, Marc Monot, Jesús L Romalde, Frédérique Le Roux

## Abstract

Phages depend on their bacterial host to replicate, but how habitat, density and diversity of the host population drive phage ecology is not well understood. Here, we addressed this question by comparing two populations of marine bacteria and their phages collected during a time series sampling in an oyster farm. *Vibrio crassostreae* reproduces more specifically in oysters. This population is genetically structured into clades of near clonal strains favoring infection by closely related phages and leading to a modular structure of the phage-bacterial infection network. *Vibrio chagasii*, on the other hand, blooms in the water column from where it can colonize oysters via filter-feeding. We found higher phage predation pressure on *V. chagasii* that did not result from a broader host range of the phages but rather from a greater burst size generating more infectious particles in the environment. We showed that contrasting patterns of genetic diversity for host and phage lead to different infection network architectures. We also provided evidence that a bloom of phages generates epigenetic and genetic variability that can be selected to counteract host defense systems.

## INTRODUCTION

Phages, the viral predator of bacteria, constitute the most abundant and diverse biological entity in the ocean and are thought to play a significant role in controlling the composition of microbial communities (Brum and Sullivan 2015, Breitbart, Bonnain et al. 2018). As obligate parasites, phages depend on their bacterial hosts to thrive in the environment and one can expect that the habitat, density and diversity of the host population are key determinants of phage ecology. Phage ecology encompasses the breadth of host strains a phage is capable of infecting (host range), the rate of intracellular phage-progeny maturation (burst) and dispersion in the ecosystem. However, the ecology of phages in marine environments remains poorly understood yet is important to understand how they control the expansion of bacterial populations in healthy natural ecosystems and to devise phage applications in disease ridden aquaculture settings (Nobrega, Costa et al. 2015, Gordillo Altamirano and Barr 2019, Kortright, Chan et al. 2019).

A prime example for mass mortalities in aquaculture caused by bacterial infections is the farming of Pacific oysters. By investigating the disease ecology of *Vibrionaceae* (here “vibrios”) in an oyster farming area, we previously observed that vibrio populations are highly dynamic (Bruto, James et al. 2017), presumably in part due to predation by phages (Piel, Bruto et al. 2022). We showed that vibrio populations shift during oyster pathogenicity events with different populations showing divergent ecological strategies. The virulent population *V. crassostreae* was abundant in diseased animals and nearly absent in the surrounding seawater, suggesting that its primary habitat and environmental reservoir is the oyster (Bruto, James et al. 2017, Piel, Bruto et al. 2020). By contrast the frequency of *V. chagasii* in oysters resembled that in the water, likely due to passive transfer by filter-feeding (Bruto, James et al. 2017). *V. chagasii* might thus first bloom in the water instead of reproducing more specifically in oysters.

We previously explored the host range of phages infecting *V. crassostreae* (Piel, Bruto et al. 2022). In a large collection of phages and their *V. crassostreae* hosts our cross infection experiments revealed large modules of interaction. Clearly delineated genomic clusters of phages (taxonomically assigned to the genus rank) were specific for distinct phylogenetic clades of *V. crassostreae*. Most of these clades contain near-clonal strains although their flexible genome can be extensive. The flexible genome encompasses a virulence plasmid that is present in a large proportion of strains (Bruto, James et al. 2017, Piel, Bruto et al. 2020) as well as other mobile genetic elements encoding for numerous phage defense systems (Piel, Bruto et al. 2022). We showed that the host range of phages results of two successive barriers: 1) the requirement for matching clade-specific bacterial receptor(s) and genus-specific phage receptor binding protein(s); 2) phage ability to overcome host defenses through epigenetic or genetic modifications (Piel, Bruto et al. 2022).

It is however unclear whether such modular structure of phage-bacteria infection network is a specific characteristic of *V. crassostreae* or a general feature resulting from vibrio-phage coevolution. Indeed another study exploring diverse populations of vibrio and their phages revealed that singleton modules of interaction were prevalent, highlighting exquisitely narrow host range of phages with respect to their local hosts in this environment (Kauffman, Chang et al. 2022). Only few larger modules were described involving 1) a broader host range *Autolykiviridae* (Kauffman, Brown et al. 2018) that infect strains from diverse species and 2) phages that infect *V. breoganii*, an algae specialist (Corzett, Elsherbini et al. 2018). This suggested that divergence in bacterial ecology and genetic diversity is reflected in interactions with different groups of phages and host range of the phages. *V. crassostreae* is confined to the oyster, increasing the encounter rates between phages and their hosts. This population is genetically structured by clones favoring infection of several hosts by a same phage. In contrast *V. chagasii* inhabits an open system, the seawater, which might be reflected in the genetic diversity of this population, the phage-predator load and/or the phage host range. Here we test these hypotheses by comparing the structure of phage-bacteria interaction networks between *V. chagasii* and *V. crassostreae*. We combine cultivation, comparative genomics, and molecular genetics to analyze a large collection of bacterial isolates and their phages. With this combination we can address how differences in the coupled host and phage ecology and diversity can structure the interaction networks and determine crucial parameters of phage ecology.

## RESULTS AND DISCUSSION

### Divergent ecology between *V. crassostreae* and *V. chagasii* populations

We previously sampled vibrios from juvenile oysters deployed in an oyster farm (Bay of Brest, France) and from the surrounding seawater on 57 dates over five months (Piel, Bruto et al. 2022). Oyster mortalities began on the 29th of May and persisted until the 25th of August (Fig.1a). A total of 194 and 252 vibrio isolates were assigned to *V. crassostreae* and *V. chagasii* respectively. *V. crassostreae* was detected only during the oyster mortality outbreak. *V. chagasii* was detected later than *V. crassostreae* and persisted after the outbreak, allowing us to assume that its role in the oyster disease is unlikely. In addition, we confirmed that this population is not preferentially associated with oyster tissues in contrast to *V. crassostreae* (Bruto, James et al. 2017).

We next compared the frequency of vibrio isolates that are susceptible to at least one phage from the same date of sampling. We used the 194 isolates of *V. crassostreae* and 252 isolates of *V. chagasii* as ‘bait’ to isolate phages from 20mL-seawater equivalents of viral concentrate (1,000X) or oyster tissues (0.1mg). Phage infection was assessed by plaque formation in soft agar overlays of host lawns mixed with the viral source. With 9% of *V. crassostreae* and 44% of *V. chagasii* isolates giving plaque(s) we found significantly fewer plaque-positive hosts for *V. crassostreae* (Fig.1b, Binomial GLM, estimate = −2.057 ± 0.278, z = −7.398, p < 0.001) indicative of a higher phage predation pressure on the *V. chagasii* population. There are several not mutually exclusive explanations for this difference: 1) *V. chagasii* phages have a broader host range and thus are able to infect more genetically diverse strains, 2) *V. chagasii* isolates have a higher susceptibility to phages because strains from this population carry less defense systems, 3) *V. chagasii* phages have a greater burst size generating more infectious particles in the environment and 4) genetic diversities of hosts and phages influence the probability to isolate co-ocurring phages.

**Figure 1.**
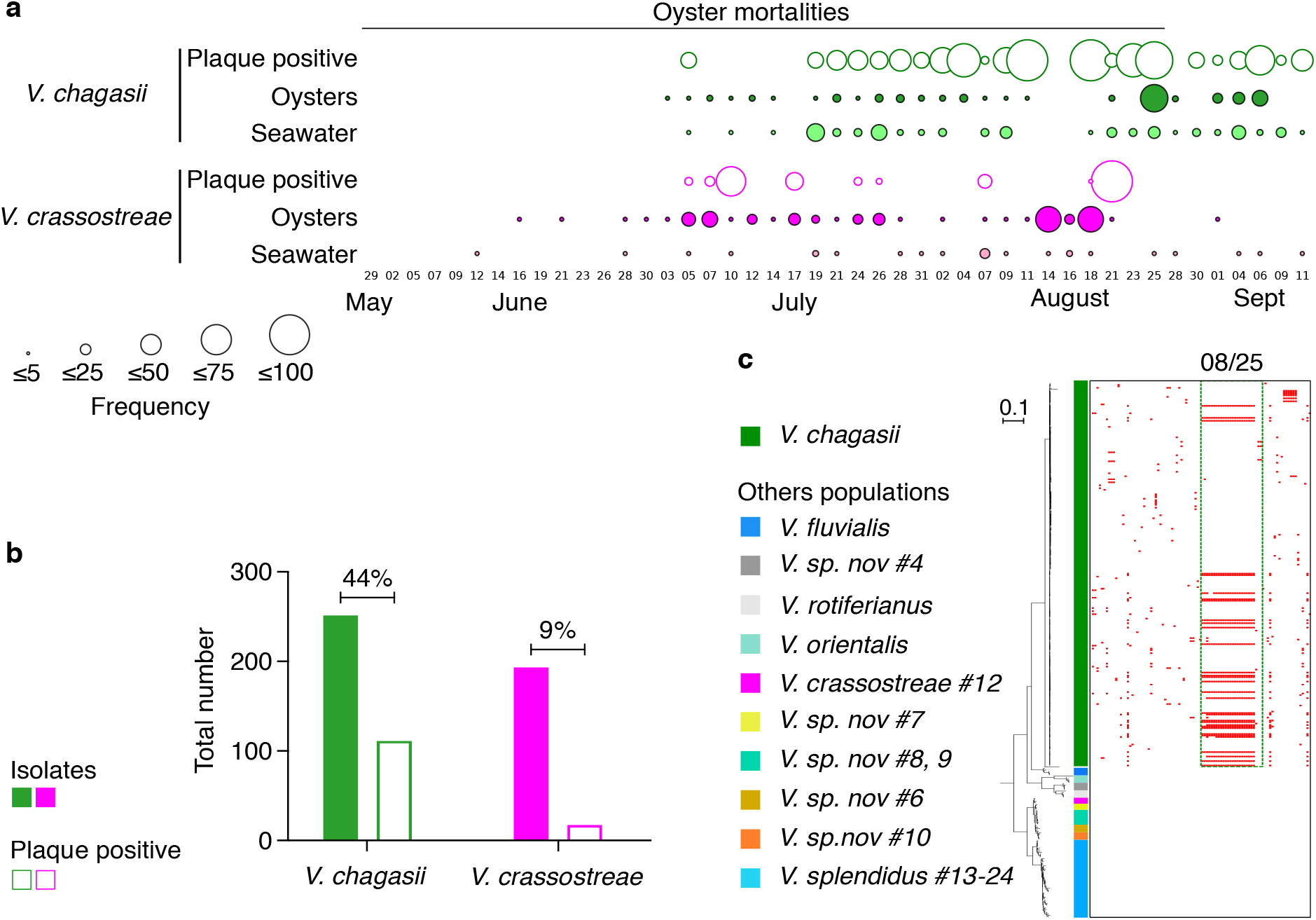
Higher predation of *V. chagasii* by highly specific phages. **a**, On each sampling date, vibrios from seawater (size fraction 1-0.2μM, light green and pink circles) or tissue from five oysters (dark green and pink circles) were selected on TCBS and genotyped to identify *V. chagasii* (green circles) or *V. crassostreae* (pink circles) isolates. The size of each circle is proportional to the frequency of *V. chagasii* or *V. crassostreae* (Freq = number of positive isolates/ randomly picked 48 colonies *100). All 252 *V. chagasii* and 194 *V. crassostreae* isolates were used as “bait” to isolate phages using viral sources from the same date. The size of each white circle is proportional to the frequency of plaque positive hosts (Freq= number of isolates from the species that give at least one PFU/total number of isolates from the species *100). **b**, The histogram indicates for each population the number of isolates and plaque positive hosts and the percentage of plaque positive hosts. **c**, Host range matrix for assay of phages on *gyrB*-sequenced hosts. Rows represent *Vibrio* strains (n=358), columns represent phages (n=95 phages isolated in Brest using *V. chagasii* as prey and ordered by date), red marks indicate infection of host. Green shade discriminates *V. chagasii* isolates from representatives of other populations (grey shades). Phages isolated on August 25 are highlight in the figure and correspond to a bloom of *V. chagasii* and infecting phages.

### *V. chagasii* infecting phages have a narrow host range

To investigate the host ranges of the phages isolated using *V. chagasii*, we purified one phage from each plaque positive host, representing a final set of 95 independent phage isolates. These phages were tested against 252 *V. chagasii* isolates from the time series and 106 representative members of other *Vibrio* species. This cross-infection assay revealed highly specific interactions between phage and bacteria since only 1047 of the 34,010 tested interactions (3%) were positive (Fig.1c) with no phages infecting members of other *Vibrio* species. Among positive interactions, 599 (57%) were obtained using phages and hosts isolated on a single date (25 August), for which we isolated the highest number of *V. chagasii* isolates (n=40/252) and their infecting phages (n=28/95). This is consistent with a role for host blooms in driving specific phage abundance as previously described (Kauffman, Chang et al. 2022). We concluded that the higher predation pressure on *V. chagasii* population does not result from a broader host range of the phages or higher susceptibility of hosts, as a similar low rate of positive interactions was obtained for *V. crassostreae*-isolated phages in cross infection assays (2.2%) (Piel, Bruto et al. 2022).

### Higher virulence of phages isolated from *V. chagasii*

The higher predation pressure on the *V. chagasii* population could alternatively be explained by a greater load of phages in their environment. In the laboratory, we observed that plaques formed by phages isolated from and infecting *V. crassostreae* are frequently smaller than those of phages isolated from and infecting *V. chagasii* (LM, estimate = −0.975 ± 0.261, t = −3.729, p < 0.001, Fig. 2a, b). We compared the titer of phages produced using *V. crassostreae* or *V. chagasii* as host (Fig. 2c) and found significant higher burst size in case of *V. chagasii* (LM, estimate = 1.374 ± 0.264, t = 5.208, p < 0.001). This suggests that a higher virulence of phages infecting *V. chagasii* results in a greater load of phages in the marine environment.

**Figure 2.**
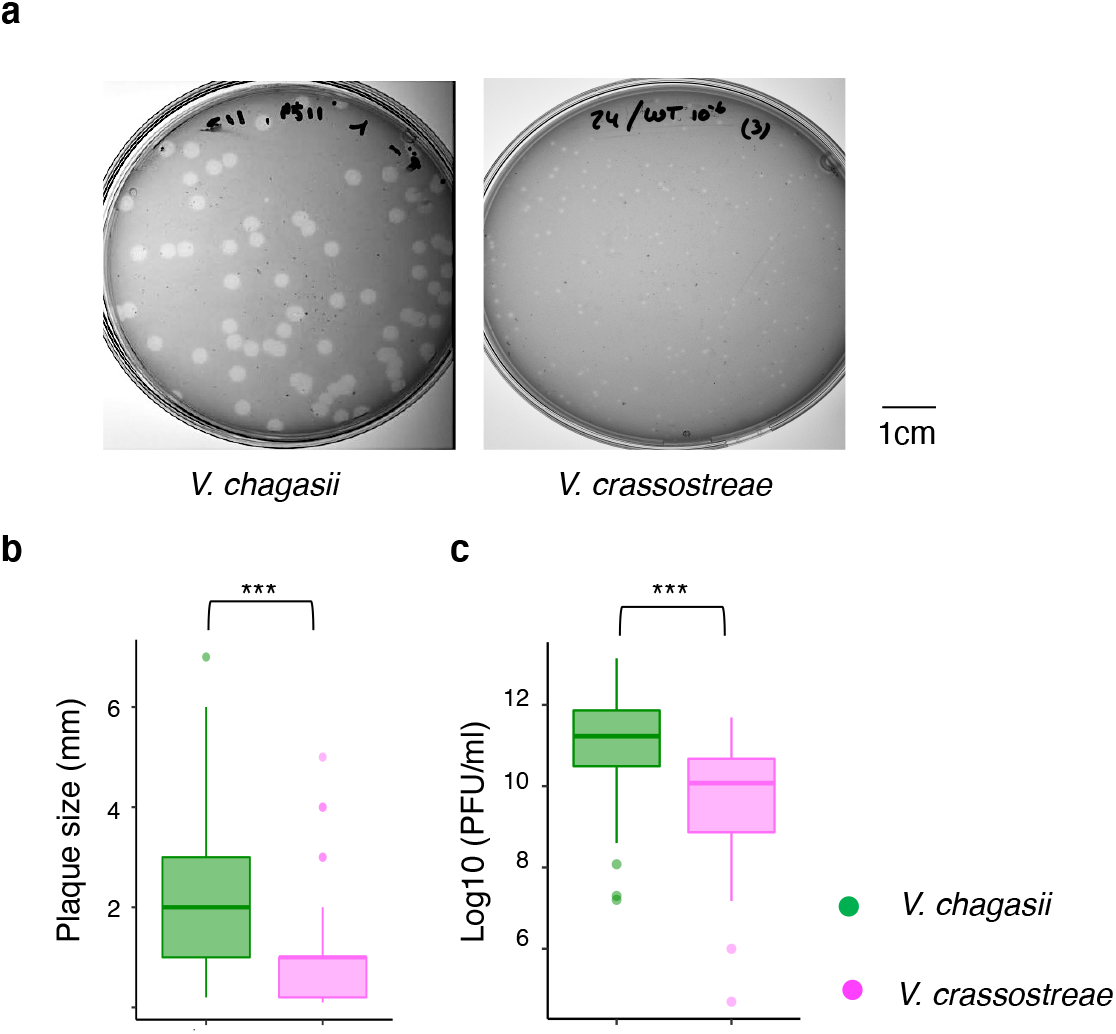
Higher virulence of phages isolated from *V. chagasii*. **a**, Typical plaques formed by phage isolated from *V. chagasii* and *V. crassostreae*. **b**, Comparison of the size of plaque for all 49 and 57 phages isolated from *V. chagasii* and *V. crassostreae* respectively revealed significant difference (LM, estimate = −0.975 ± 0.261, t = −3.729, p < 0.001). **c**, The titer of phages obtained from high titer stock preparation using phages infecting *V. chagasii* or *V. crassostreae* was used as a proxy to show a significant higher burst size for phages infecting *V. chagasii* (LM, estimate = −1.374 ± 0.264, t = −5.208, p < 0.001).

### Genome analysis reveals differences in population structure and ecological traits

In *V. crassostreae* we showed that phage infection depends initially on their ability to adsorb to the host, which restricts their host range to a specific bacterial clade. Consequently, large modules observed in the infection network resulted from the particular genetic structure of this population in clades (Fig. 3a). To analyze how *V. chagasii’*s genetic structure explains the phage-bacteria interaction matrix, we sequenced and assembled the genomes of selected strains (Table S1). A total of 136 representatives were chosen by clustering the *V. chagasii* isolates with identical *gyrB* sequences and identical patterns of susceptibility in the cross-infection assays. The core genome phylogeny and pairwise ANI values revealed striking differences with *V. crassostreae* (Fig. 3a). In *V. crassostreae* the genetic distance is low within clades and large between clades (Fig. 3b). In *V. chagasii*, genetic distances within the species are more homogenous, lower overall but larger than the within-clade distance in *V. crassostreae*.

**Figure 3.**
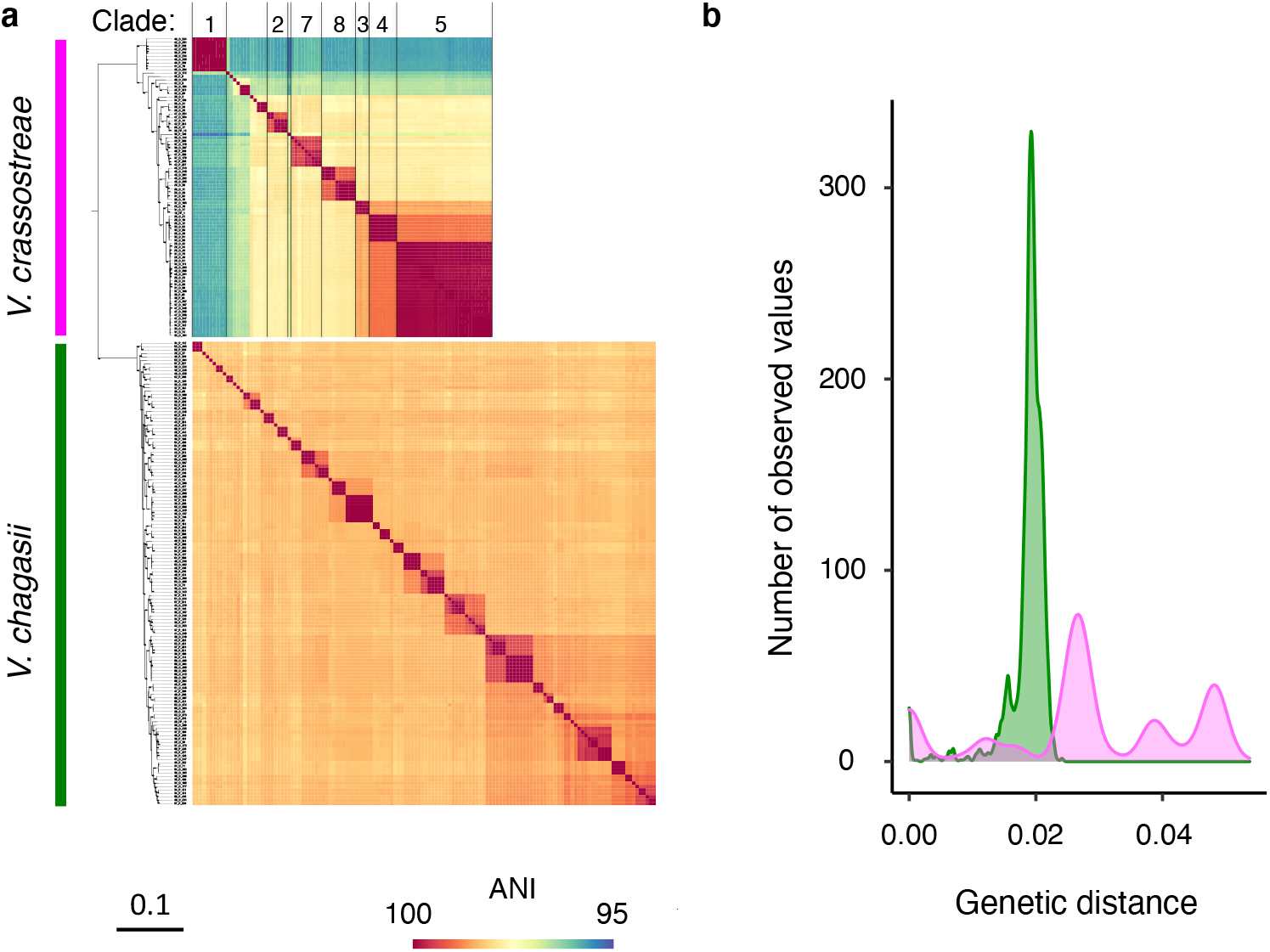
Genome diversity of *V. crassostreae*. and *V. chagasii*. **a**, Core genome phylogeny based on 2689 gene families of 88 *V. crassostreae* and 136 *V. chagasii* isolates from the time series and pairwise ANI values revealing highly structured clades in *V. crassostreae* but not in *V. chagasii*. The clade numbers refer to (Piel, Bruto et al. 2022). **b**, The resulting distribution of pairwise genetic distance within *V. chagasii* (green) and *V. crassostreae* (pink) showed striking difference between these species (Two-sample Kolmogorov-Smirnov test, D = 0.762, *p* < 0.001). *V. chagasii* shows a unimodal distribution with few clonal strains and a homogenous pairwise genetic distances with a maximum differentiation of 0.024, whereas *V. crassostreae* shows a multimodal distribution corresponding to the distinct clades with low within-clade and large between clade genetic distance.

Genomes of *V. chagasii* vary in size between 5.3 to 6.5 Mb (Table S1), which is indicative of an extensive flexible genome diversity among these closely related isolates. Recent work has shown that most phage defense systems are encoded on mobile genetic elements (Hussain, Dubert et al. 2021). In *V. crassostreae* we showed that the number of defense systems is strongly correlated with bacterial genome size and both measures correlate with the number of different infecting phages (i.e. host susceptibility) (Piel, Bruto et al. 2022). Although we found a wealth of different phage defense systems in *V. chagasii* as well (Table S2), the number of these systems was not correlated with the genome size and host resistance (number of defense systems versus genome size, Spearman rho = −0.03, p =0.66; number of defense systems versus number of infecting phages per strain; Spearman rho = −0.06, p = 0.4862).

A total of 191 genes were exclusively found in all *V. chagasii* isolates compared to only 53 private genes in *V. crassostreae* isolates (Table S3). Among the *V. chagasii* specific genes we found clusters presumably involved in bacteria attachment (tight adherence and curling) and motility (flagellar) which might indicate a broader colonization potential of *V. chagasii* population involving more different habitats. On the other hand, none of the *V. chagasii* strains carries the virulence plasmid pGV1512 that is present in 72% of *V. crassostreae* isolates. This plasmid encodes a type 6 secretion system cytotoxic for oyster immune cells (Piel, Bruto et al. 2020). Hence population specific genes seem congruent with their respective lifestyle, generalism for *V. chagasii* and host specificity for *V. crassostreae* and we can also expect that the difference in the structure of these populations influences the diversity of phages that infect them.

### Phages isolated from *V. chagasii* are more diverse

To analyze the diversity of phages isolated using *V. chagasii* we grouped them according to their host, date of isolation and patterns of infectivity leading to 49 representatives. We investigated phage morphology, genome sequence, and taxonomy. Electron microscopy revealed that all phages belong to the *Caudovirales*, including myoviruses (n=16), podoviruses (n=22) and siphoviruses (n=11) (Fig.S1). Genome sequencing of these double-stranded DNA viruses revealed extensive variability in genome length (from 31,648 to 161,442 bp) and gene content (45 to 305 predicted coding sequences) (Table S4). To explore the genome diversity of these phages we used the Virus Intergenomic Distance Calculator (VIRIDIC, (Moraru, Varsani et al. 2020)). This led us to group the 49 phages infecting *V. chagasii* into 25 VIRIDIC genera (ANI>70%) and 41 species (ANI>95%) (Fig. 4a). By comparison, the 57 phages isolated using *V. crassostreae* were assigned to 19 genera and 23 species (Fig. 4b), indicating that phages isolated from *V. chagasii* are genetically more diverse particularly on the species level.

**Figure 4.**
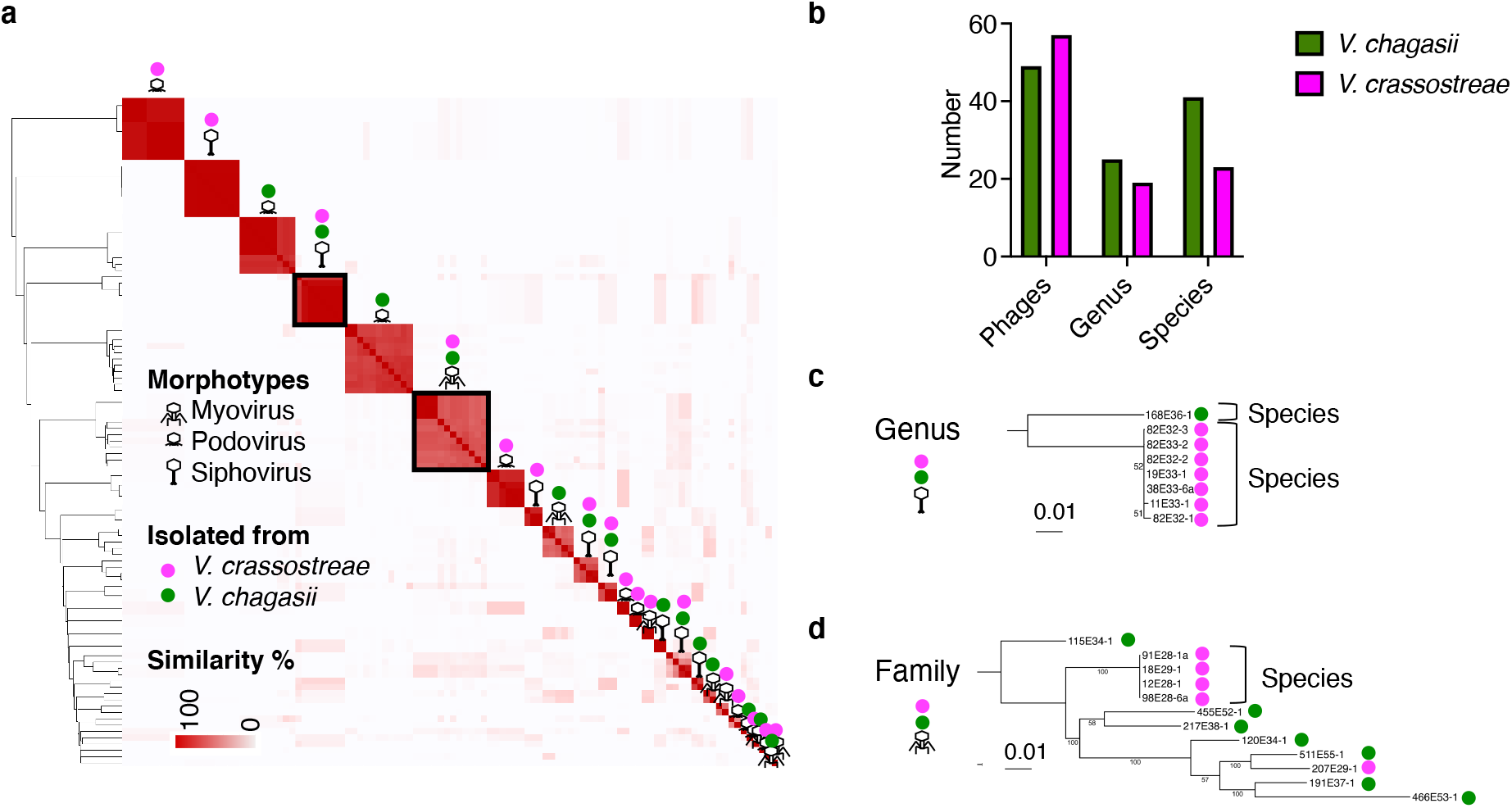
Diversity of phages isolated using *V. chagasii* and *V. crassostreae* as host. **a**, VIRIDIC intergenomic similarity between phage genomes. The 106 phages included in the study were grouped into 43 clusters assigned to VIRIDIC genus rank (>70% identities). **b**, The 49 and 57 phages infecting *V. chagasii* and *V. crassostreae* were assigned to 25 and 18 genera respectively. Phage morphotypes are indicated by specific icons for siphoviruses (long tail), myoviruses (medium tail) and podoviruses (short tail) (see Fig. S1, for TEM). The species of the host used to isolate the phage are indicated by a pink and green circle for *V. crassostreae* and *V. chagasii* respectively. **c**, Only one genus (siphoviruses framed in bold in a) contains phages isolated using *V. crassostreae* or *V. chagasii*. This genus splits into two VIRIDIC species (>95% identities), one is representing by a single phage isolated from *V. chagasii* and the second is containing seven phages all isolated from *V. crassostreae*. The phylogenic tree is based on 41 core proteins (25% identities and 80% coverage) from the genus of siphoviruses isolated from *V. chagasii* or *V. crassostreae*. **d**, Some genera were grouped into clusters assigned to VIRIDIC family rank (>50% identities). Among these families, five grouped genera isolated from *V. chagasii* or *V. crassostreae* (e.g. the myoviruses framed in bold in a). The phylogenic tree is based on 35 core proteins (25% identities and 80% coverage) from the family of myoviruses isolated from *V. chagasii* or *V. crassostreae*.

Only one genus (Genus 1, Podoviruses, Table S4) contained 11 phages predicted as temperate phages by BACPHILIP (score 0.975). A gene encoding a tyrosine recombinase was detected in these genomes. However tyrosine recombinase constitutes a large family involved in a wide variety of biological processes (post-replicative segregation, genetic switches and movement of mobile genetic elements including phages) (Smyshlyaev, Bateman et al. 2021). Yet, functional annotation of these proteins is still limited by their large diversity and lack of experimental data. Finally, this prediction was contradicted by the absence of integration of the phages in the *V. chagasii* genomes.

We identified a single genus (Genus 10, Siphoviruses, Table S4) that contains phages infecting both vibrio species (Fig. 4c). This genus splits into two VIRIDIC species (identities >95%), the first one containing a single phage isolated from *V. chagasii* and the second one containing seven phages isolated from *V. crassostreae*. Some genera of phages isolated from *V. crassostreae* or *V. chagasii* could be clustered in VIRIDIC families (identities >50%). A core proteome phylogeny of the best represented family indicated that phages differentiated from a common ancestor into genus and species in their respective vibrio host species (Fig. 4d). Interestingly, the phylogeny also revealed that phage jumped from one host species to another. We propose that vibrio diversity and containment in oyster could expand the genomic diversification of the coevolving viral population, promoting phage host jumps, as previously described for the gut microbia (De Sordi, Khanna et al. 2017).

### The structure of the infection network is driven by the phylogenetic distances within **the** host population

The contrasting patterns of genetic diversity for hosts and phages between the two *Vibrio* species could potentially also cause the observed differences in the structure of the infection network (Fig. 5a). In *V. chagasii*, the size of module was significantly smaller (negative binomial GLM, estimate = 1.234 ± 0.241, z = 5.121, p < 0.001, Fig. 5b). These modules involve genetically related hosts killed by genetically related phages. Phages isolated using *V. chagasii* are more diverse than those from *V. crassostreae* which might explain at least partially the difference in module size. Smaller modules might also suggest a higher specialization of the phages, i.e. a lower number of susceptible hosts. Comparing pairwise number of shared phages and phylogenetic distance we found that for both species the closer the vibrio strains are related the more phages they share (Fig. 5c). While both species showed a bimodal distribution of sharing probabilities, it is noteworthy that highest likelihood to share phages was at genetic distances close to 0 for *V. chagasii* whereas the distribution was shifted upward with maximal probabilities at 0.01 for *V. crassostreae* (Fig. 5c). This supports the higher degree of specialization of *V. chagasii* phages. In fact, only one phage (382E49-1) out of 49 was able to infect strains with larger genetic distances (>0.024). We therefore conclude that smaller modules in *V. chagasii* are consequence of a lower number of hosts with low genetic distance as well as a higher diversity of the phages and a higher degree of specialization. Overall, these data highlight the importance of considering genetic structures of hosts and phages when interpreting phage bacteria networks.

**Figure 5.**
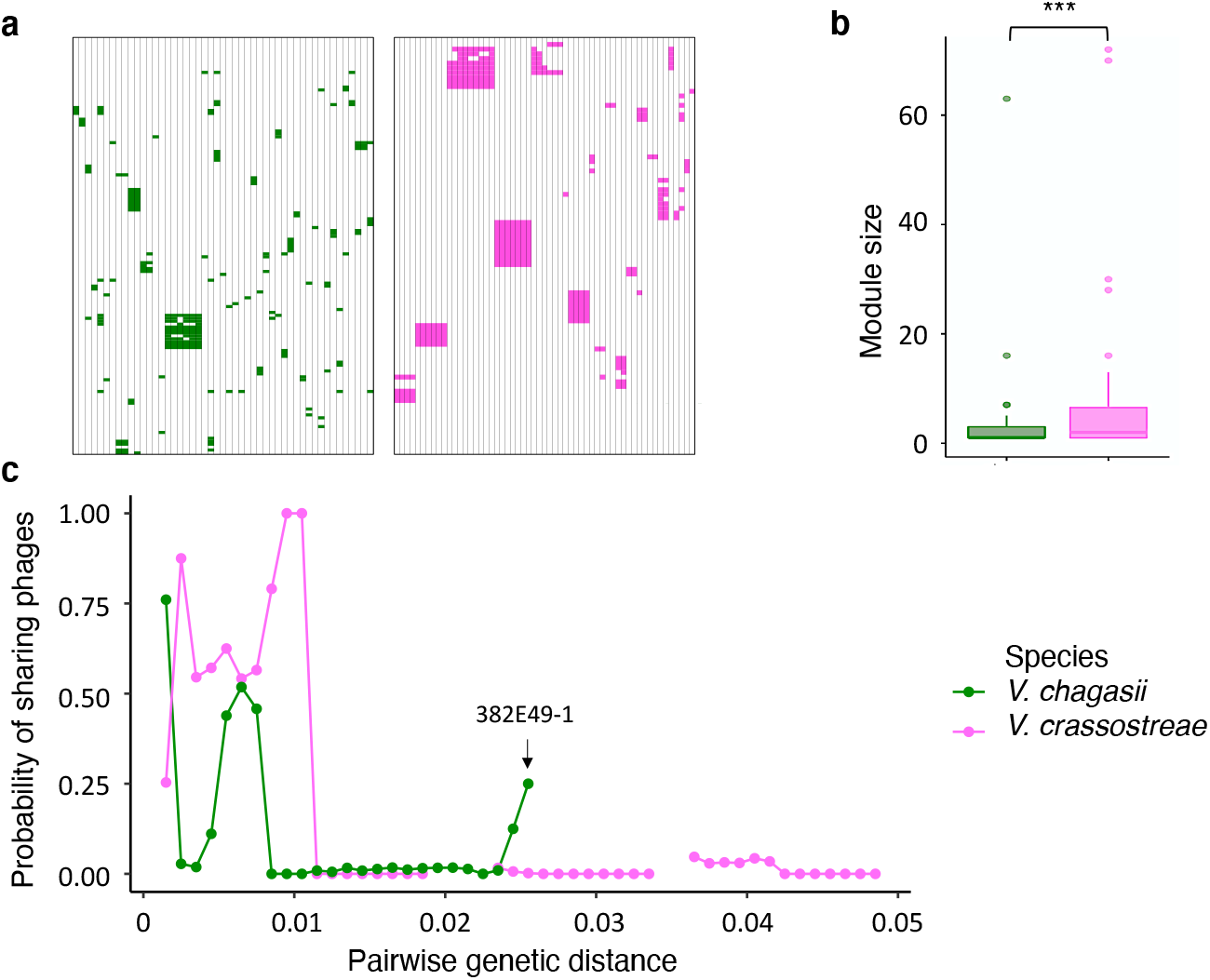
Modularity is driven by phylogenetic distances within host population. **a**, Rows represent sequenced *Vibrio* strains ordered according to the Maximum likelihood core genome phylogeny of *V. chagasii* (n=126; green PBIN) and *V. crassostreae* strains (n=88; pink PBIN) (see the phylogeny Fig. 3a). Columns represent sequenced phages (49 and 57 isolated from *V. chagasii* and *V. crassostreae*) ordered by VIRIDIC genus (see the dendrogram Fig. 4a). Colored squares indicate killing of the respective vibrio isolate by the phage. **b**, The size of modules was significantly different between *V. chagasii* and *V. crassostreae* (negative binomial GLM, estimate = 1.234 ± 0.241, z = 5.121, p < 0.001). **c**, Sharing probability (i.e. proportion of phages shared in all pairs within a window of genetic distances) as a function of host genetic distance. Sliding window with an interval size of 0.003 and a step size of 0.001.

### Bloom of phages generate large phage numbers, epigenetic and genetic variability

Having an exquisitely narrow host range, phages have to find strategies that increase the encounter rates with their host. Being an oyster specialist, a higher density of *V. crassostreae* in animal tissues might favor physical contact with their phages. We showed that phages infecting *V. chagasii* have a higher burst presumably increasing the load of phages in open seawater. In addition host blooms can drive increase in abundances of specific phage types as observed in the time series the 25^th^ August (Fig. 1a and c). Such blooms might also generate phage genetic diversity. Indeed among the phages with sequenced genomes isolated on this date, five belong to the same species (species 26, genus 18, podoviruses, Table S4) and differed by a unique single nucleotide polymorphism (SNP) (Fig. S2). We compared the virulence of these phages for diverse susceptible hosts using efficiency of plating (EOP), i.e. the titer of the phage on a given bacteria compared to the titer on the strain used to isolate the phage (original host) (Fig.6a). This revealed that only the phage 409E50-1 showed noticeable variation separating four out of 12 vibrio strains with low susceptibility. These four “LS” (low susceptibility) strains showed significantly lower susceptibility than the eight remaining strains “HS” (high susceptibility) (LM, estimate = −3.322 ± 0.081, t = −40.609, p < 0.001 Fig.6b). Lower infection could not be attributed to difference in cell-surface phage receptors, as the phage 409E50-1 adsorbs to all tested isolates regardless of the production of progeny (Fig. S3). We thus concluded that an intracellular defense system found in LS strains controls the efficiency of phage 409E50-1 infection while other phages from the species are able to counteract this defense.

Genome comparisons revealed 276 genes present in all LS and absent in all HS strains (Table S5). Among them we found two known phage defense systems, the phosphorothioation-sensing bacterial defense system (sspBCDE) targetting the viral DNA (Xiong, Wu et al. 2020) and the cyclic-oligonucleotide-based anti-phage signalling systems (CBASS_I) leading to host suicide (Millman, Melamed et al. 2020). However the double deletion of these systems in 36_P_168 did not change the susceptibility of the host (not shown). The LS specific defense mechanism(s) acting on 409E50-1 thus still remains to be identified.

We also sought to understand how the other phages of this species escape the LS defense. A phage can escape from host defenses by epigenetic or genetic modifications. The phages 409E50-1 and 521E56-1 diverge by their original host strain (respectively 50_O_409 and 56_P_521) and a single SNP in their genome. To explore the hypothesis of epigenetic modification, we produced these phages in the two host strains (Fig. 6c). When the phage 409E50-1 was propagated on vibrio 56_P_521 its infectivity for LS strains was strongly increased compared to the progeny obtained from the original host (LMM, estimate = 2.016 ± 0.296, t = 6.821, p = 0.002). The infectivity of phage 521E56-1 was slightly increased when produced on 50_O_409 compared to its original host (Fig. 6c, LMM estimate = 0.656 ± 0.062, t = 10.62, p < 0.001). However, the effect estimate of 409E50-1was three times larger than the one for 521E56-1 indicating that the latter was less affected by the host used to produce progenies. This suggests that the vibrio strain 56_P_521 confers an epigenetic advantage to phage 409E50-1 and we hypothesized that the phage 521E56-1 does not require host-mediated modification because of genetic divergence.

**Figure 6:**
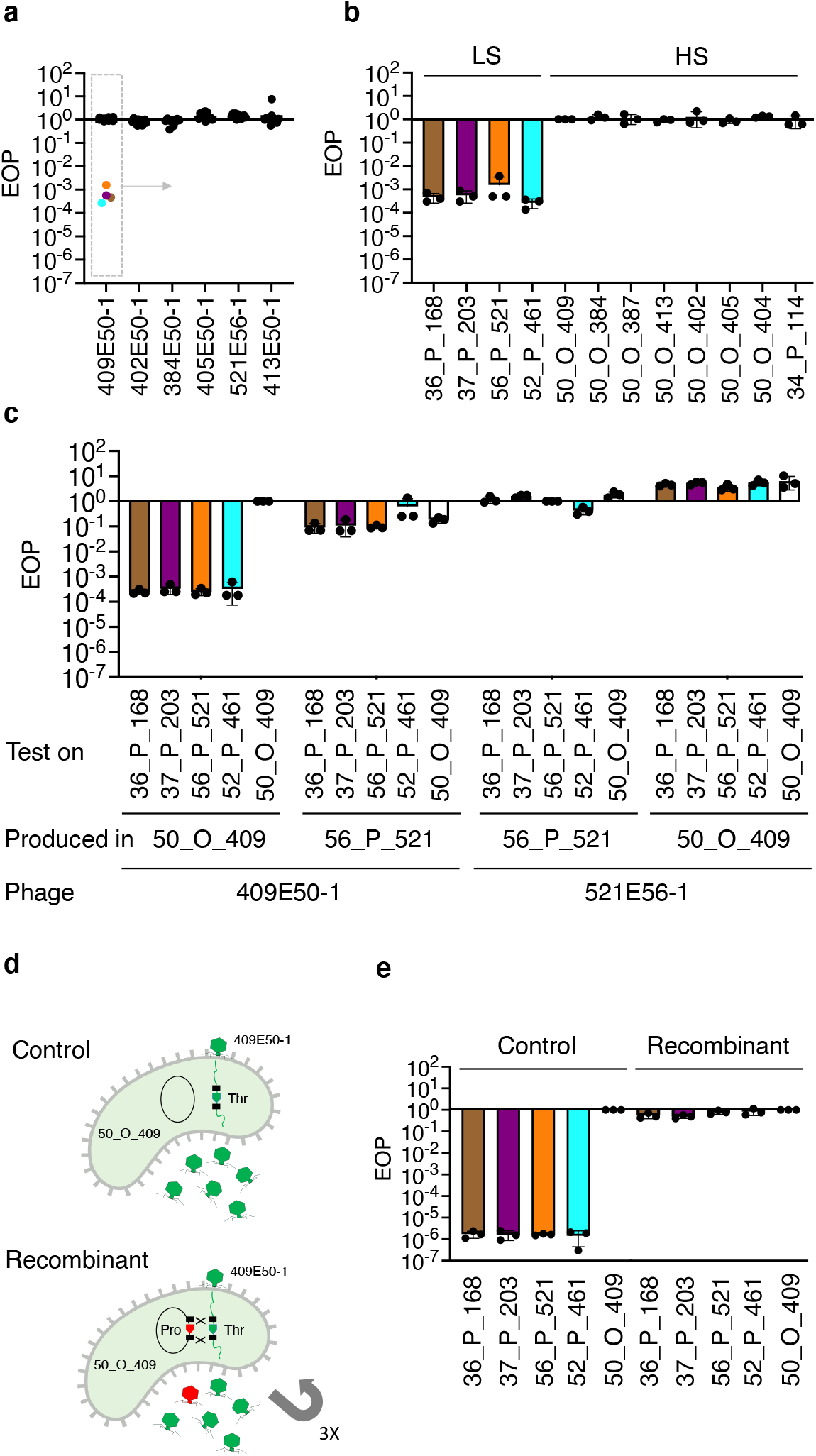
Epigenetic and genetic modification are involved in phage adaptation. **a**, Efficiency of plating (EOP, the titer of the phage on a given bacteria divided by the titer on the strain used to isolate the phage or ‘original host’) of six phages from VIRIDIC species 26 (Fig. S2) on 12 *V. chagasii* strains. Individual dots correspond to the mean from three independent experiments. **b**, EOP of the phage 409E50-1 infecting *V. chagasii* strains with low susceptibility (LS) or high susceptibility (HS). Bar charts show the mean +/-s.d. from three independent experiments (individual dots). **c**, EOP obtained using *V. chagasii* strains 50_O_409 (HS) or 56_P_521 (LS) to produce the phages 409E50-1 or 521E56-1. Results were standardized by experiment, i.e. the production and testing on original host gave 1 for each experiment. **d**, Graphic summary of the strategy used to exchange the nucleotide T to G and codon Thr to Pro in phage 409E50-1. **e**, EOP of the recombinant phages or control infecting the four *V. chagasii* strains with low susceptibility (LS). Results were standardized by experiment, i.e. the production in original host (50_0_409) gave 1 for each experiment.

The phage 409E50-1 differs from 521E56-1 by a single SNP localized in a gene encoding a protein of unknown function (label VP495E501_p0076 in phage 409E50-1; 125 amino acid). This SNP results in the change of a threonine (409E50-1) by a proline (521E56-1) at the amino acid position 8 (Fig. S4). We exchanged the allele T=>G in the phage 409E50-1 by recombination (see method and Fig. 6d). We noticed a significant decrease in the infectivity of the control phages for LS strains (EOP of 10^−6^ instead of 10^−4^ for the ancestral phage, Fig. 6b and 6c), the origin of this loss of infectivity remaining to be elucidated. Irrespective of this difference, control phages infections showed significantly lower EOP on LS strains compared to the original host, 50_0_409 (LM, estimate = −5.843 ± 0.123, t = −47.571, p < 0.001). The recombinant phages (T=>G) were on average 10^6^ fold more infectious than control phages on LS strains and did not differ significantly from the original host, 50_0_409 (Fig. 6e, LM, estimate = −0.160 ± 0.120, t = −1.327, p = 0.196). We can therefore show that a single SNP provides protection from the unknown phage defense system present in LS strains. We speculate that this locus is also the target for epigenetic protection confered by the host 56_P_521 to the phage 409E50-1.

HS strains were isolated from oysters collected on August 25. The absence of strain specific genes and a low number of SNPs (2 to 25) confirms their clonality (Fig. S5). By comparison, LS strains were more diverse (5 to 823 SNPs, 40 genes specific to strain 52_P_461 and 10 genes specific to strain 37_P_203) and were isolated from seawater collected before and after 25^th^ of August. The bloom like proliferation observed on the 25^th^ of August therefore most likely resulted from the clonal expansion of a HS strain in oyster(s). This expansion of a *V. chagasii* clone in oysters coincides with the disappearance of *V. crassostreae* in the animals. This could be explained by biotic or abiotic factors that are no longer conducive to the replication of *V. crassostreae* in the oyster, leaving this niche accessible to another population (population shift). This specific *V. chagasii* clone might also outcompete *V. crassostreae* which is already threatened by phages. Finally, the clonal expansion presumably drove the overproportional increase of phages assigned to species 26. It is conceivable that this large phage population generated genetic by mutation, including the previously identified SNP that lead to adaptation to a host defense system. These adapted phages were in turn able to reproduce in LS strains after the HS bloom.

## CONCLUSION

In this study we explored how host ecology and genetic structure feeds back on phage ecology (host range, burst size and population dynamics) and genetic diversity of both host and phages. While *V. crassostreae* is an oyster specialist, *V. chagasii* has a generalist lifestyle that coincides with a greater potential for mobility and colonization. Divergence in bacterial ecology is reflected by difference in host phage infection networks resulting in different predation pressures. The higher predation pressure on the *V. chagasii* population does not result from a broader host range of the phages or higher susceptibility of host but rather from a greater burst size generating more infectious particles in the environment. We speculate that phages with a higher burst are selectively favored in the seawater column because the host encounter rates are lower in this open system than in oysters.

Contrasting patterns of genetic diversity for host and phage between the species result in different structures of the respective phage bacteria interaction networks. In *V. crassostreae*, large modules of interaction result from the particular genetic structure of this population in clades of near clonal strains favoring the isolation of closely related phages. Smaller modules in *V. chagasii* are consequence of a lower number of clonal hosts. More diverse hosts used as prey will lead to the isolation of more diverse phages. A higher specialization of the phage infecting *V. chagasii* was however supported by the highest likelihood to share phages when genetic distances are small. Host blooms result in a high number of phages and we showed that coupled with the high burst size a high number of phages can generate adaptive genetic variability. Selection can work on this increased variability and may favor variants that can also infect host outside the bloom, that were less susceptible to the original phage. The more generalist lifestyle of *V. chagasii* with blooms in- and outside of oyster hosts may thus increase genetic variability of phages, but prevent the formation of modules in a host with low population structure.

## MATERIAL AND METHODS

### Sampling

Sampling was performed at the same time as for Piel et al 2022. Briefly, samples were collected from an oyster farm located at the Bay of Brest (Pointe du Château, 48° 20′ 06.19′′ N, 4° 19′ 06.37′′ W), every Monday, Wednesday and Friday from the 3^rd^ of May to the 11^th^ of September 2017. Specific Pathogen Free (SPF) juvenile oysters (Petton, Bruto et al. 2015)were deployed in the field in batches of 100 animals. When the first mortalities were observed in the first batch, another batch of SPF animals was placed in the field, leading to the consecutive deployment of seven batches from the 26^th^ of April to the 11^th^ of September. Oyster mortalities were recorded on each sampling day. Seawater temperature reached 16°C on the 22^nd^ of May, a previously observed threshold for oyster mortalities (de Lorgeril, Lucasson et al. 2018). Mortalities began on the 29^th^ of May and persisted until the 25^th^ of August. On each sampling date, five living oysters were collected from a batch showing less than 50% mortalities. The animals were cleaned, shucked, weighed and 2216 Marine Broth (MB) was added (10 mg/ml) for homogenization using an ultra-turrax. A volume of 100 μL homogenate was used for vibrio isolation, the remaining volume was centrifuged (10 min,17,000 g), the supernatant filtered through a 0.2 μm filter and stored at 4°C until the phage isolation stage. Two liters of seawater were collected and size fractionated as previously described (Bruto, James et al. 2017). Bacterial cells from 0.2 μm filters were suspended in 2 mL MB and 100 μL of this suspension was used for vibrio isolation. The iron chloride flocculation method was used to generate 1000-fold concentrated viral samples from 2 liters passaged through a 0.2 μm filter, following the previously described protocol (Kauffman, Brown et al. 2018). Virus-flocculates were suspended in 2mL 0.1M EDTA, 0.2M MgCl_2_, 0.2M oxalate buffer at pH6 and stored at 4°C until the phage isolation stage.

### *Vibrio* isolation, identification and genome analysis

#### Isolation and identification

Vibrios from seawater or oyster tissues were selected on thiosulfate-citrate-bile salts-sucrose agar (TCBS). Roughly 48 colonies were randomly picked from each plate and re-isolated once on TCBS, then on 2216 Marine agar (MA). Colonies were screened by PCR targeting the *r5*.*2* gene encoding for a regulator (Lemire, Goudenege et al. 2014). PCR positive isolates were grown in MB and stored at −80°C in 10% DMSO. Bacteria were grown overnight in 5mL MB and DNA extracted using an extraction kit (Wizard, Promega) according to the manufacturer’s instructions. Taxonomic assignment was further refined by *gyrB* gene sequencing to identify *V. crassostreae* and *V. chagasii* isolates. The partial *gyrB* gene was amplified using degenerate primers (Table S6), Sanger sequenced (Macrogen) were manually corrected with the chromatogram. Sequences were aligned with Muscle and phylogenetic reconstruction was done with RAxML version 8 GTR model of evolution, a gamma model and default parameters (Stamatakis 2006).

#### Genome sequencing, assembly and annotation

*V. chagasii* libraries were prepared from 500 ng of genomic DNA using MGIEasy Universal DNA Library Prep Set following the manufacturer’s instruction. The library pool were circularised and sequenced using DNBSEQ-G400 (BGI) 100bp paired-end by the Biomics platform at the Pasteur Institute. Reads trimming was done using Trimmomatic v0.39 (LEADING:3, TRAILING:3, SLIDINGWINDOW:4:15, MINLEN:36) (Bolger, Lohse et al. 2014). *De novo* reads assembly was performed using Spades version 3.15.2 (--careful --cov-cutoff auto -k 21,33,55,77 -m 10). In order to confirm BGI’s sequencing quality, we compare these assembly results with those obtained with for 20 same *V. chagasii* strains sequenced by Illumina HiSeq using QUAST v5.0.2 (Gurevich, Saveliev et al. 2013) (Table S7). We confirmed the previous benchmark studies showing similar sequencing and assembly qualities across both platform (Foox, Tighe et al. 2021).

Computational prediction of coding sequences and functional assignments were performed using the automated annotation pipeline implemented in the MicroScope platform (Vallenet, Belda et al. 2013). Phage defense genes annotation was performed using Defense-Finder version 1.0.8 (Tesson, Herve et al. 2022) with MacSyFinder models version 1.1.0 and default options. Persistent genome phylogeny was done using the PanACoTA workflow (Perrin and Rocha 2021) version 1.3.1-dev2 with mash (version 2.3) distance filtering from 0 to 0.3, prodigal version 2.6.3 for syntaxic annotation, mmseqs version 2.11.e1a1c for genes clustering, mafft version 7.407 for persistent genome alignment defined as 90.0% of all genomes in each family and IQ-TREE (Nguyen, Schmidt et al. 2015)version 2.0.3 based on GTR model and 1000 bootstrap for tree construction.

#### Comparative genomics

The average nucleotide identity (ANI) value of genomes was determined using fastANI version 1.32 (fragment length 500) (Jain, Rodriguez et al. 2018). *Vibrio crassostreae* and *Vibrio chagasii* core, variable, accessory and specific genomes was established using mmseqs2 reciprocal best hit version 13-45111 with 80% identities on 80% coverage thresholds (Steinegger and Soding 2018).

### Phage isolation, identification and genome analysis

#### Phage isolation

We used the 194 and 252 isolates of *V. crassostreae* and *V. chagasii* as ‘bait’ to isolate phages from the same date of sampling from 20mL-seawater equivalents of viral concentrate (1,000X) or oyster tissues (0.1mg). Phage infection was assessed by plaque formation in soft agar overlays of host lawns mixed with a viral source. We obtained plaque(s) for 9% and 58% of *V. crassostreae* and *V. chagasii* isolates respectively (Fig.1b). Concerning *V. chagasii*, we then purified one phage from each plaque positive host, representing a final set of 95 phage isolates. In addition, we selected 57 *V. crassostreae*-phages isolated from the same date of sampling as their host (Piel, Bruto et al. 2022). The size of the plaques was measured for each phage isolated from *V. chagasii* and *V. crassostreae*. To generate high titer stocks, plaque plugs were first eluted in 500 μl of MB for 24 hours at 4°C, 0.2-μm filtered to remove bacteria, and re-isolated three times on the sensitive host for purification before storage at 4°C and, after supplementation of 25% glycerol at −80°C. High titer stocks (>10^9^ PFU/ml) were generated by confluent lysis in agar overlays and phage concentration was determined drop spots of ten-fold serial dilutions onto bacterial host lawns.

#### Electron microscopy

Following concentration on centrifugal filtration devices (Millipore, amicon Ultra centrifugal filter, Ultracel 30K, UFC903024), 20 μl of the phage concentrate were adsorbed for 10 min to a formvar film on a carbon-coated 300 mesh copper grid (FF-300 Cu formvar square mesh Cu, delta microscopy). The adsorbed samples were negatively contrasted with 2% Uranyl acetate (EMS, Hatfield, PA, USA). Imaging was performed using a Jeol JEM-1400 Transmission Electron Microscope equipped with an Orious Gatan camera at the platform MERIMAGE (Station Biologique, Roscoff, France).

#### DNA extraction

Phage suspensions (10 mL, >10^10^ PFU/mL) were concentrated to approximately 500μL on centrifugal filtration devices (30 kDa Millipore Ultra Centrifugal Filter, Ultracel UFC903024) and washed with 1/100 MB to decrease salt concentration. The concentrated phages were next treated for 30 min at 37°C with 10 μL of DNAse (Promega) and 2.5μL of RNAse (Macherey-Nagel) at 1000 unit and 3.5 mg/mL, respectively. The nucleases were inactivated by adding EDTA (20 mM, pH8). DNA extractions encompassed a first step of protein lysis (0.02 M EDTA pH 8.0, 0.5 mg/mL proteinase K, 0.5% SDS) for 30 min incubation at 55°C, a phenol chloroform extraction and an ethanol precipitation. DNA was visualized by agarose gel electrophoresis (0.7% agarose, 50V, overnight at 4°C) and quantified using QuBit.

#### Genome sequencing, assembly

Phages were sequenced by the Biomics platform at the Pasteur Institute. The sequencing library was prepared using a TruSeq Illumina kit, 2 × 75 bp paired end, and sequenced on a NextSeq550 Illumina sequencer. After reads trimming done with Trimmomatic v0.39 (LEADING:3, TRAILING:3, SLIDINGWINDOW:4:15, MINLEN:36) (Bolger, Lohse et al. 2014), de novo assembly was perfomed using SPAdes v3.15.2 (--careful --cov-cutoff auto -k 21,33,55,77 -m 10). Vector contamination contigs were excluded using the UniVec Database and one-contig phage genome was manually decircularized. The syntaxic annotation was done with PHANOTATE v1.5.0 (McNair, Zhou et al. 2019). Large terminase subunit was identified using DIAMOND v2.0.8.146 (blastp, default parameters) (Buchfink, Reuter et al. 2021) on previous annotated vibrio large terminase and using HMMSCAN (HMMER v3.3.2, default parameters with evalue≤10^−3^) with PFAM profiles (PF03354.17, PF04466.15, PF05876.14, PF06056.14) (Mistry, Chuguransky et al. 2021). The phage genome was manually ordered starting from the gene for the large terminase subunit.

#### Genome annotation

The tRNA were identified with tRNAscan-SE v2.0.9 (Chan and Lowe 2019). The functional annotation used multiple approaches. First we used HMMSCAN (evalue≤10^−3^) with PVOGS profiles (Grazziotin, Koonin et al. 2017). We also used DIAMOND blastP against first the Nahant collection genomes and after against others bacterial RefSeq (version 212) and phage GenBank genomes (2022-24-06, with 30% identities and 50% coverage) (Kauffman, Brown et al. 2018, Buchfink, Reuter et al. 2021). Finally we added InterProScan v5.52-86.0 (evalue≤10^−3^) and eggNOG v5.0.2 annotation results (Jones, Binns et al. 2014, Huerta-Cepas, Szklarczyk et al. 2019).

#### Clustering, lifestyle and comparative genomic

We clustered phage using VIRIDIC v1.0r3.6 (default parameters) (Moraru, Varsani et al. 2020). Intergenomic similarities were identified using BLASTN pairwise comparisons. Viruses assignment into family (**≥**50% similarities) genera (**≥**70% similarities) and species (**≥**95% similarities) ranks follows the International Committee on Taxonomy of Viruses (ICTV) genome identity thresholds. Phage lifestyle was predicted using BACPHILIP version 0.9.3-alpha and lysogeny associated genes was searched using InterProScan (v.5.52-86.0) based on integrase (IPR002104), resolvase (IPR006119), replicative transposases (IPR004189), and transcriptional repressor domains (IPR010982, IPR010744, IPR001387, IPR032499). Absence of prophage integration was also checked using BLASTn searches between both phage and bacterial genomes (evalue ≤0.01) (Hussain, Dubert et al. 2021). All comparative genomic analysis was done using MMseqs2 version 13-45111 for protein clustering (Steinegger and Soding 2018), FAMSA version 1.6.2 for genes/proteins alignments (Deorowicz, Debudaj-Grabysz et al. 2016) and IQ-TREE version 2.1.2 for phylogenetic tree construction (Nguyen, Schmidt et al. 2015).

### Host range determination

#### Single-phage-by-single-host host range infection assay

Host range assays were carried out using an electronic multichannel pipette by spotting 5 μL of the phage suspension normalized at 2×10^5^ PFU/ml (10^3^ PFU/spot) on the agar overlay inoculated with the tested host. Plates were incubated overnight at RT and plaque formation was observed after 24 hours. Concerning *V. chagasii*, spot assays were performed, first by infecting 358 vibrio strains (252 *V. chagasii* + 106 strains belonging to other vibrio populations) with 95 phages in duplicate. Then, the host range of 49 selected phages was confirmed on 136 *V. chagasii* hosts. Concerning *V. crassostreae*, results were obtained in (Piel, Bruto et al. 2022).

#### Efficiency of plating (EOP)

Ten-fold serial dilutions of phages from a high titer stock (>10^9^ PFU/mL) were prepared and 10 μL of each dilution were pipetted onto bacterial host lawns. The EOP was calculated as the ratio between the titer of a phage on a given strain by the titer of the same phage on its reference strain (host used to isolate and produce the phage).

#### Adsorption estimation

Phage adsorption experiments were performed as previously described (Hyman and Abedon 2009). Phages were mixed with exponentially growing cells (OD 0.3; 10^7^ CFU/mL) at a MOI of 0.01 and incubated at RT without agitation. At different time points, 250 μL of the culture was transferred in a 1.5 mL tube containing 50 μL of chloroform and centrifuged at 17,000g for 5 min. The supernatant was 10-fold serially diluted and drop spotted onto a fresh lawn of a sensitive host to quantify the remaining free phage particles. In this assay, a drop in the number of infectious particles at 15 or 30 min indicates bacteriophage adsorption.

### Statistical analyses

Statistical analyses were based on linear models (LM) when data were normally distributed or on negative bionmial generalized linear models (negbin GLM) when data were left skewed. Correlations were based on non-parametric Spearman’s rank correlations. Paired tests were analysed as linear mixed models adding the paired factor as a random intercept. All analyses were performed using the R statistical environment (v.4.0.5).

### Molecular microbiology

#### Strains and plasmids

All plasmids and strains used or constructed in the present study are described in Table S8 and S9. *V. chagasii* isolates were grown in Marine broth (MA) or LB+0.5 M NaCl at RT. *Escherichia coli* strains were grown in LB at 37°C. Chloramphenicol (Cm; 5 or 25μg/mL for *V. chagasii* and *E. coli*, respectively), thymidine (0.3 mM) and diaminopimelate (0.3 mM) were added as supplements when necessary. Induction of the P_BAD_ promoter was achieved by the addition of 0.2% L-arabinose to the growth media, and conversely, was repressed by the addition of 1% D-glucose. Conjugation between *E. coli* and *V. chagasii* were performed at 30°C as described previously (Le Roux, Binesse et al. 2007).

#### Clonings

For vibrio knock out, all clonings in the suicide vector pSW7848T were performed using Herculase II fusion DNA polymerase (Agilent) for PCR amplification and the Gibson Assembly Master Mix (New England Biolabs, NEB) according to the manufacturer instructions. For recombination of the phage 409E50-1, a region of 227 bp flanking the SNP (T in 409E50-1, G in other phages from species 26) was amplified using the Herculase II and DNA fom the phage 521E56-1 and cloned by Gibson Assembly in pMRB (Le Roux, Davis et al. 2011) instead of the *gfp* gene. All clonings were first confirmed by digesting plasmid minipreps with specific restriction enzymes and second by sequencing the insert (Eurofins).

#### Vibrio mutagenesis

Knock out was performed by cloning 500bp fragments flanking the region in the pSW7848T (Le Roux, Binesse et al. 2007). This suicide vector encodes the *ccdB* toxin gene under the control of an arabinose-inducible and glucose-repressible promoter, *P*_*BAD*_. Selection of the plasmid-borne drug marker on Cm and glucose resulted from integration of pSW7848T in the genome. The second recombination leading to pSW7848T elimination was selected on arabinose-containing media.

#### Phage mutagenesis

Recombinant phage (T=>G in p0076 of phage 409E50-1) was engineered using double crossing over with a plasmid carrying a 227 region of homology to the phage genome 521E56-1 (Fig. 6d). This plasmid, or as control the original pMRB-*gfp*, was transferred by conjugation to the strain 50_O_409. Plate lysates were generated by mixing 100 μL of an overnight culture of the transconjugant with the 409E50-1 phage and plating in 5 mL agar overlay. After the development of a confluent lysis of lawns, the lysate was harvested by addition of 7 mL of MB, shredding of the agar overlay and stored ON at 4°C for diffusion of phage particles. The lysates were next centrifuged, the supernatant filtered through 0.2 μm filter and stored at 4°C. We could not enriched the recombinant phages by infecting LS strains because these strains confer epigenetic protection to the 409E50-1 phage. We then decided to perform several successive passages of the phage on the strain carrying the plasmid, testing the infectivity of the phages on the original strain or on an LS strain at each passage. After three passages, a different titer between the putative recombinant and controls was observed. The p0076 gene was PCR amplified using single plaque as template (eight recombinant or eight control phages) and primers flanking the 227 region in the phage genome (external primers). The SNP was confirmed by sequencing. Three phages control or recombinant were tested by EOP on four LS strains in addition to the source 50_O_409.

## Data availability

Sequenced genomes have been deposited under the ENA Project with accession numbers PRJEB53320 for *V. chagasii* and PRJEB53960 for their infecting phages.

## Materials availability

All vibrio strains, phage strains and plasmids are available upon request.

## ACKNOWLEDGEMENTS

We warmly thank Sylvain Gandon, François Blanquart and Martin Polz for fruitful discussions on the manuscript. We thank Sabine Chenivesse, Sophie Le Panse, the staff of the station Ifremer Argenton and Bouin, the ABIMS (Roscoff) and LABGeM (Evry) platforms for technical assistance. We thank all members of the “GV team” for support with field sampling. We thank Zachary Allouche, Biomics Platform, C2RT, Institut Pasteur, Paris, France, supported by France Génomique (ANR-10-INBS-09) and IBISA. This work was supported by fundings from the European Research Council (ERC) under the European Union’s Horizon 2020 research and innovation program (grant agreement No 884988, Advanced ERC Dynamic) and the Agence Nationale de la Recherche (ANR-20-CE35-0014 « RESISTE ») to FLR. R.B.-C. acknowledges the Spanish Ministerio de Ciencia e Innovación for his FPI predoctoral contract (BES-2017-079730).

## SUPPLEMENTARY DATA

**Figure S1:**
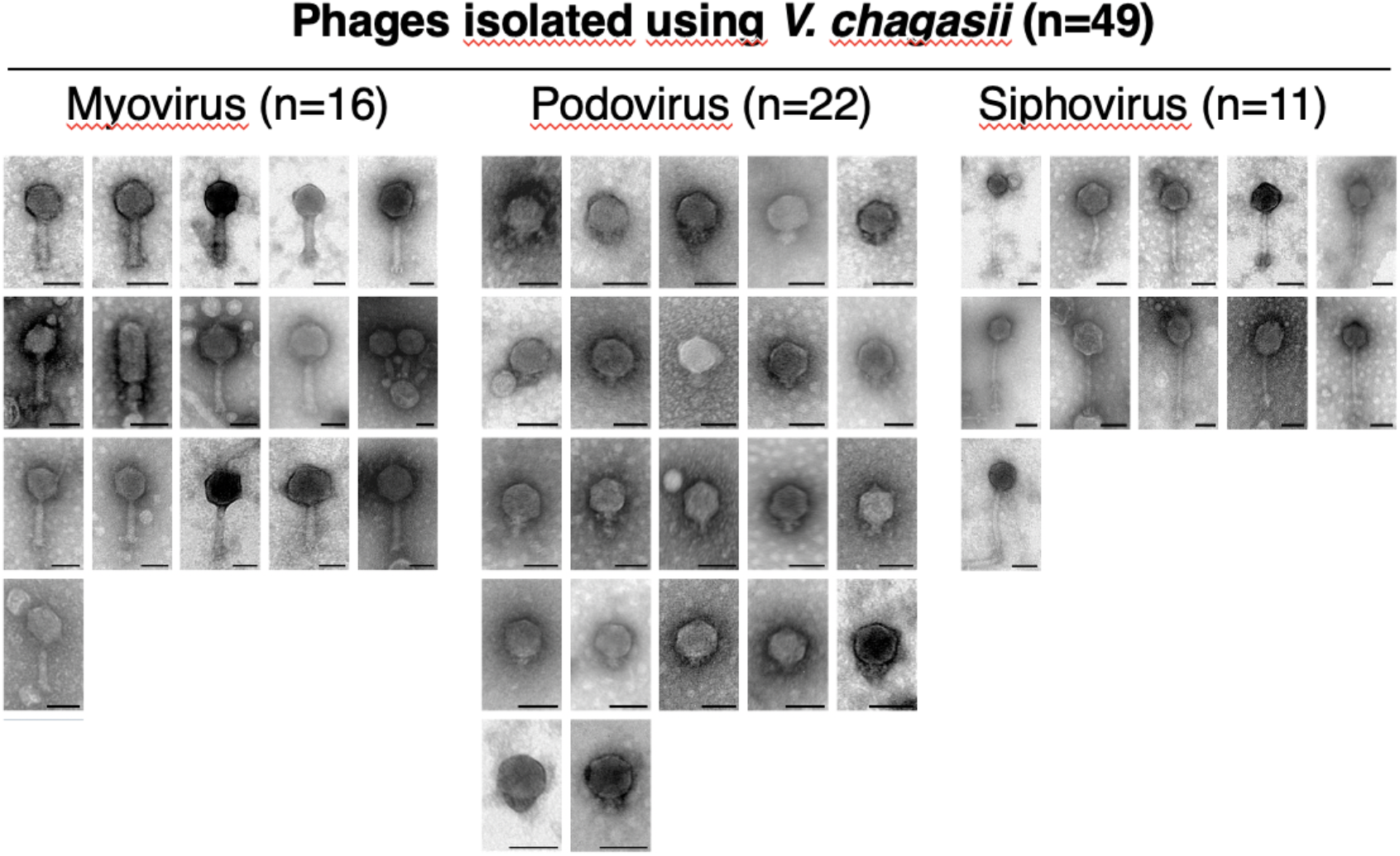
Transmission electron micrographs revealed the 49 phages belong to the *Caudovirales*, with 16 myoviruses, 22 podoviruses and 11 siphoviruses. Scale bars represent 50nm.

**Figure S2:**
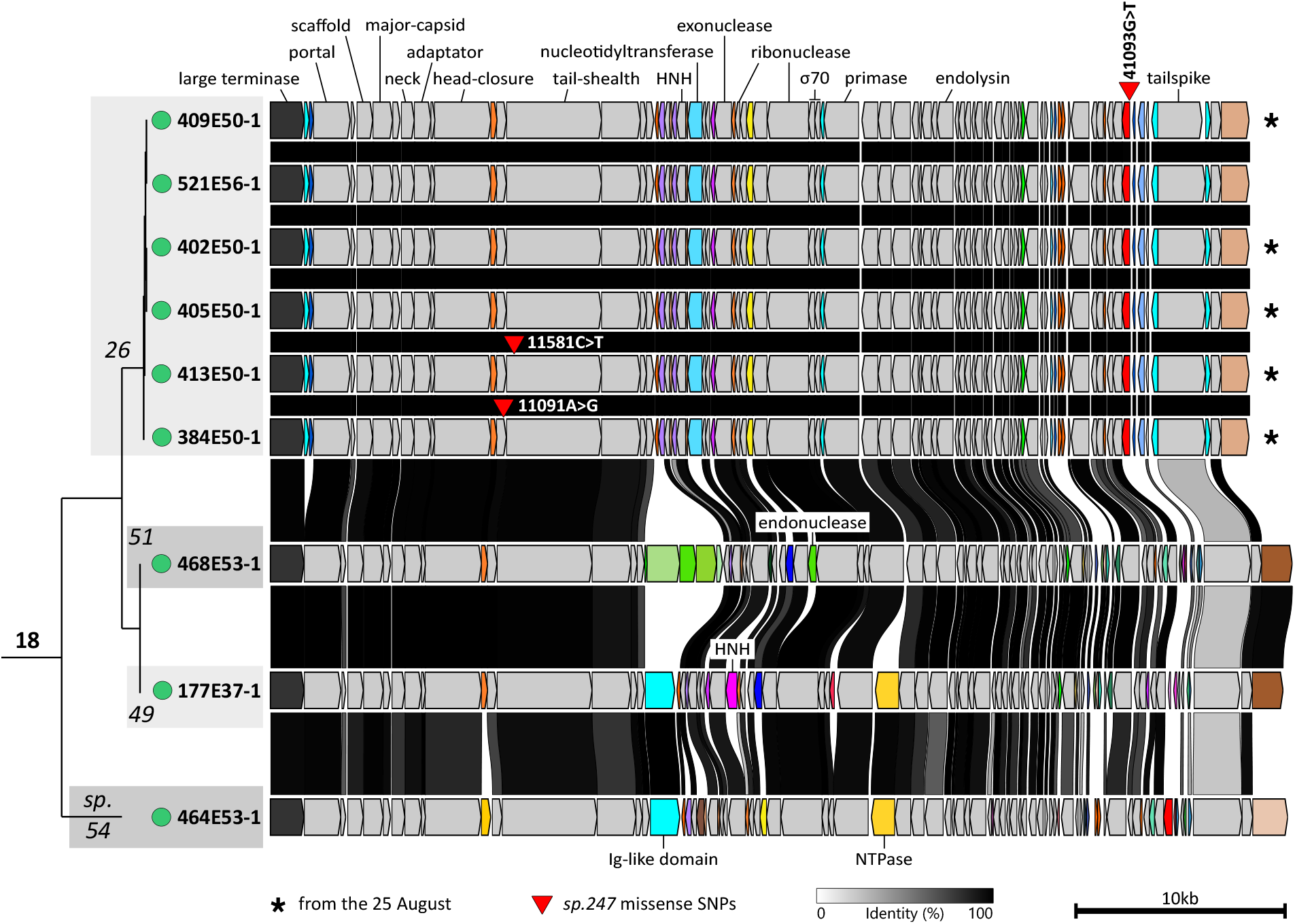
Core phylogenetic tree and genes synteny for phages from the genus 18. Core proteins are indicated in grey (with 30% identity and 80% coverage thresholds), large terminase subunit in black and accessory proteins in colors. The tree was built using IQ-TREE2 with 1000 bootstraps and the GTR model. Six phages (5 isolated the 25 August, indicated with *) are clustered in the species 26. Phage genotypes differ by a single SNP. The phage 409E50-1 shows a G to T mutation in the gene VP495E501_p0076 encoding for an unknown function.

**Figure S3:**
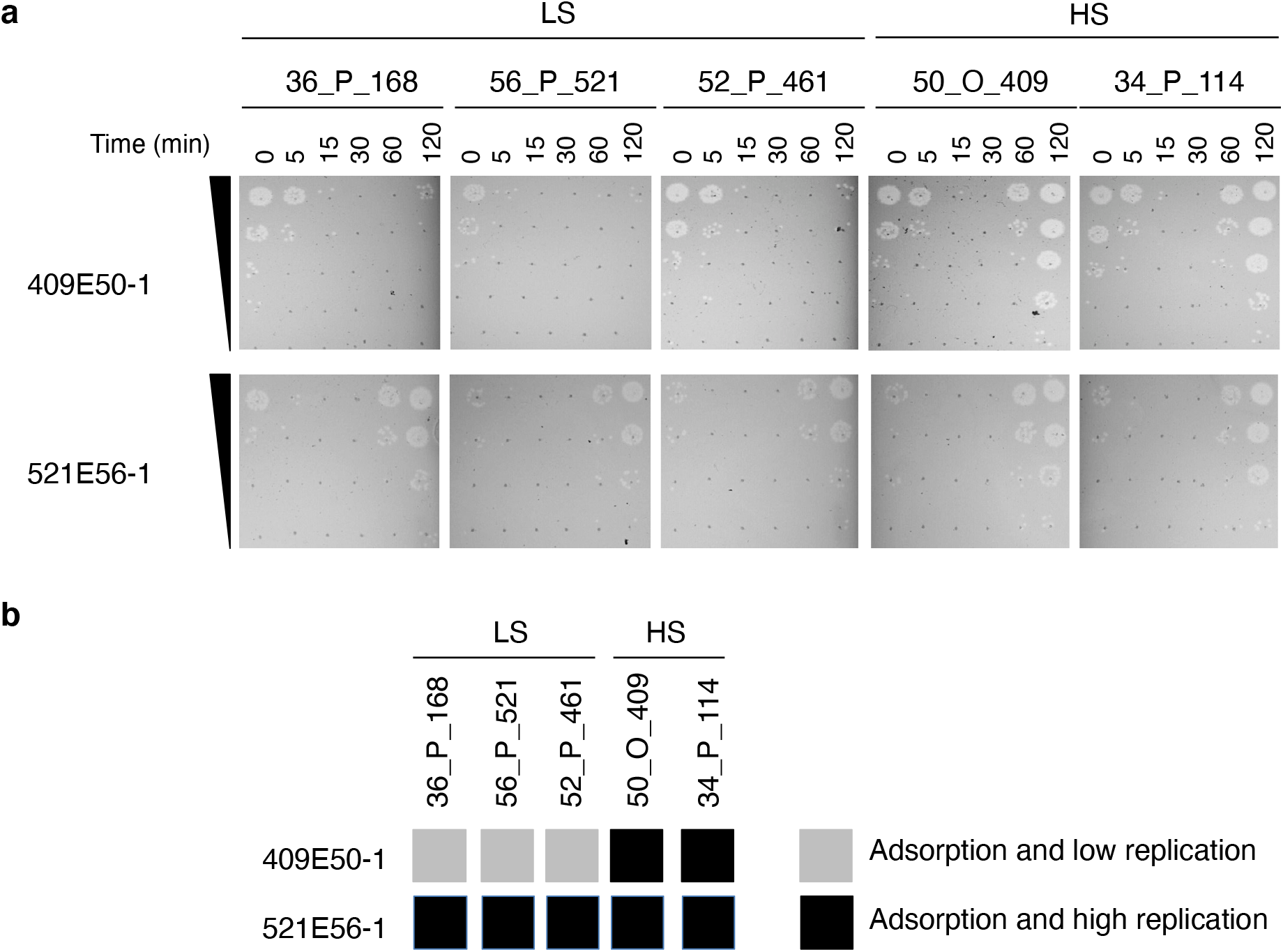
**a**, Adsorption assays of 409E50-1 and 521E56-1 on three LS strains and two HS strains. Both phages can adsorb to all tested isolates regardless of the efficacity of replication. **b**, Graphic summary of the results.

**Figure S4:**
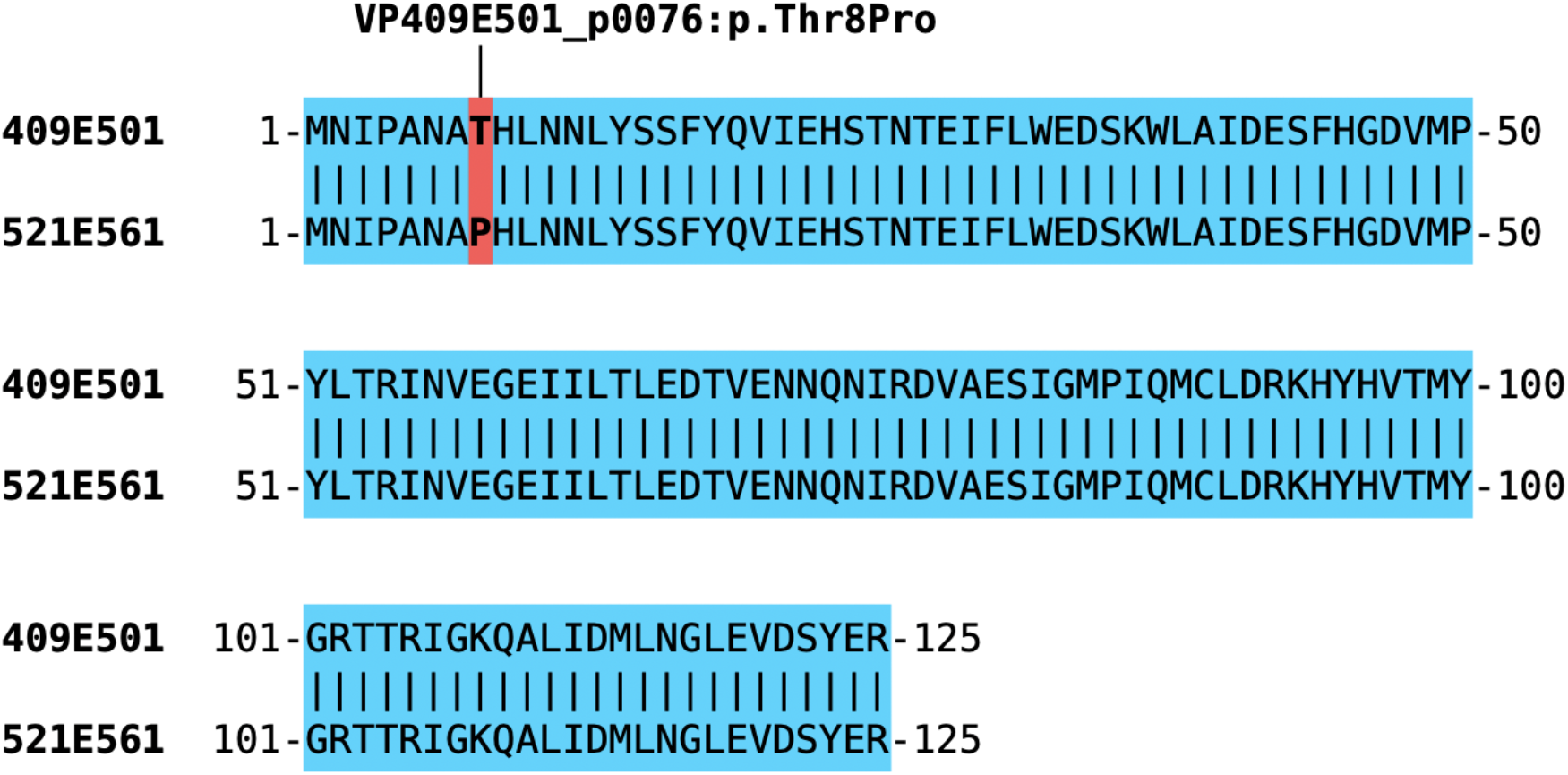
SNP results in the change of a threonine (409E50-1) by a proline (521E56-1) at the amino acid position 8 in the gene labelled p0076 from 409E50-1 and encoding for unknown function.

**Figure S5:**
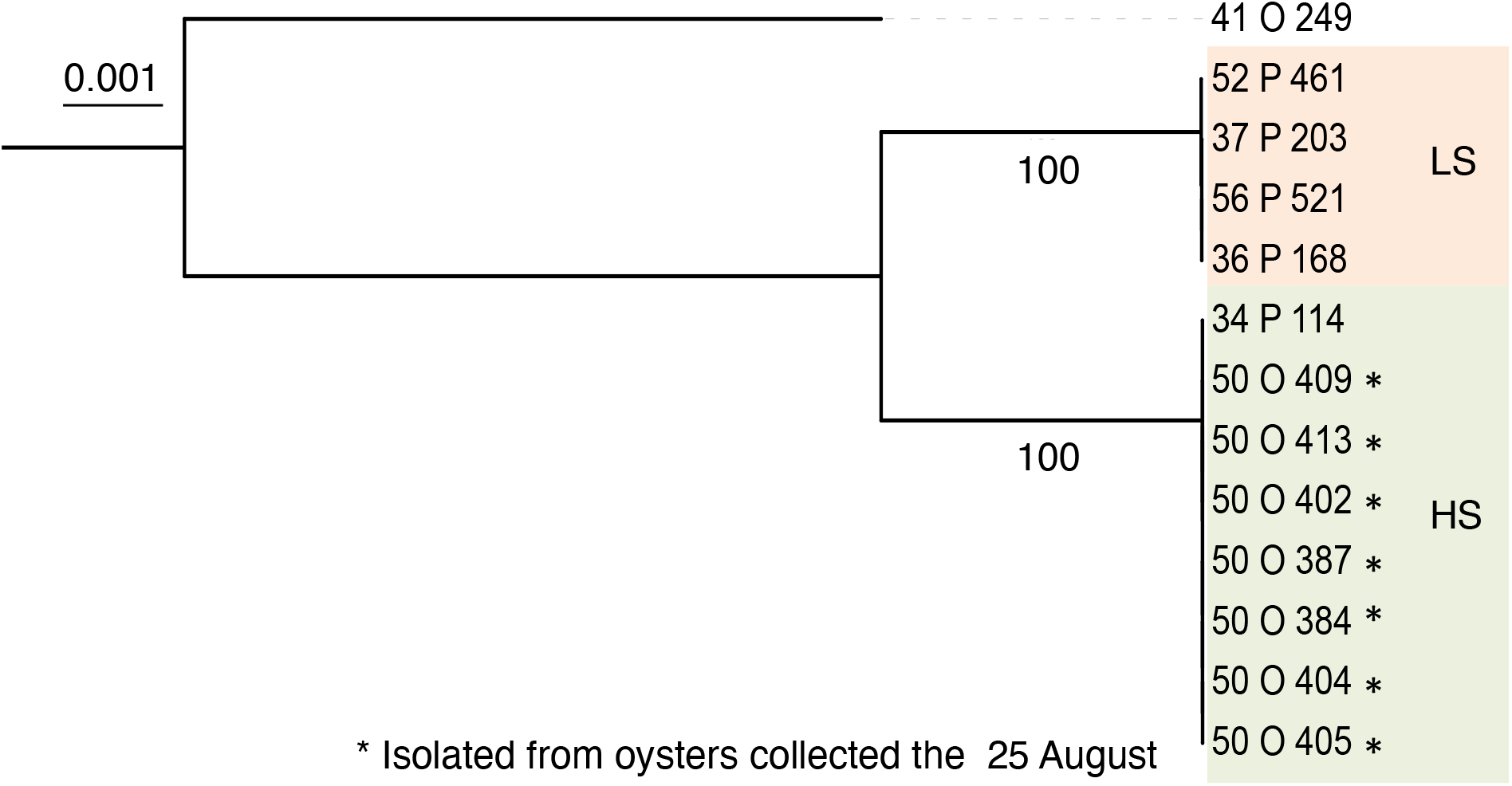
Phylogenetic analysis based on concatenated alignments of nucleic acid sequences of 4494 persistent genes (present at 80% identity and 80% coverage in 90% of all genomes) from 12 *V. chagasii* strains from the killing module. The *V. chagasii* strain 41_O_249 was used as an outgroup. The tree was built by the Maximum-Likelihood method with a GTR model based on a sequence alignment generated by mafft. Branch lengths are drawn to scale and are proportional to the number of nucleotide changes. Numbers at each node represent the percentage value given by bootstrap analysis of 1000 replicates.

